# Distribution and abundance of tetraether lipid cyclization genes in terrestrial hot springs reflects pH

**DOI:** 10.1101/2022.08.15.504015

**Authors:** Laura N. Blum, Daniel R. Colman, Emiley A. Eloe-Fadrosh, Matthew Kellom, Eric S. Boyd, Olga Zhaxybayeva, William D. Leavitt

## Abstract

Many Archaea produce membrane-spanning glycerol dibiphytanyl glycerol tetraether (GDGTs) lipids that serve as unique biomarkers of past environments. These lipids can contain up to eight cyclopentane rings, where an increase in ring cyclization is generally associated with growth in more acidic, higher temperature, or more energy limited conditions. Recently the genes that encode GDGT ring synthases, *grsAB*, were identified and characterized in model thermoacidophiles *Sulfolobus acidocaldarius* and *Saccharolobus solfataricus*. However, the distribution and abundance of *grs* homologs across environments inhabited by these and related Archaea remains unknown. To address this, we examined the distribution of *grs* homologs in archaeal and bacterial cultivar genomes, single cell genomes, metagenomes, and metatranscriptomes from thermal springs across the planet, where temperature, pH, and geochemical data take at time of sampling. The relative abundance of *grs* in these microbial communities exhibits a strong negative correlation with pH, and weak positive correlation with temperature. Genomes and metagenome-assembled genomes (MAGs) from Archaea that encode two or more copies of *grs* are significantly more widespread in low pH springs. Homologs of *grs* were detected in MAGs from 12 archaeal classes, with the most well-represented being the Thermoproteia. Homologs of *grs* were also detected among several classes of uncultured Archaea, including the Korarchaeia, Bathyarchaeia, and Hadarchaeia. Several Nitrososphaeria MAGs had high copy numbers of *grs* (> 3), and the functional role of these copies cannot yet be explained. Notably, *grs* genes were also found in MAGs from the bacterial class Acidobacteria. Based on phylogenetic analyses, it is likely that Acidobacteria acquired these genes horizontally from Archaea. Broadly, our results highlight the key role of *grs*-catalyzed lipid cyclization in the diversification of Archaea in hot and acidic environments.

## 1.0 Introduction

Microorganisms on Earth today provide insight into life’s evolutionary responses to environmental changes over geologic time. Microbes from the domain Archaea are well-adapted to survive in environments characterized by multiple extremes, such as high temperature and acidic pH, which together impose chronic energy limitation on cells (1–3). Notable among the evolutionary innovations that enable Archaea to thrive in extreme environments are their isoprenoid tetraether membrane lipids (4–7). One prevalent class of archaeal membrane lipids, the glycerol dibiphytanyl glycerol tetraethers (GDGTs), consist of isoprenoid building blocks that form monolayer cell membranes comprising 40 carbon (C40) chains. The addition of cyclopentane rings to the hydrophobic core of GDGTs is an important adaptation of many Archaea to extremes in temperature, pH, and energy availability (4,8–12). The addition of rings alters the lipids’ tertiary structure such that the membrane packs together more tightly (4,6–8,13). This structural modification increases the rigidity and decreases the permeability of the membrane, enabling the cell to remain intact at higher temperatures and preventing the influx of protons at acidic pH (7,14–19).

Archaeal GDGTs contain between zero and eight cyclopentane rings within the two hydrophobic core C40 isoprenoid chains, with zero to four rings per chain (20–22). In some strains, GDGTs can also contain a single cyclohexyl ring, in the lipid known as crenarchaeol, which to-date is only found within the Nitrososphaerota (syn. Thaumarchaeota) (21–24). Culture studies have shown that Archaea systematically increase the extent of GDGT cyclization in response to decreases in pH and increases in temperature (4,8,10,25–29). More recent culture studies have shown that the extent of GDGT cyclization is also increased under electron donor- or acceptor-limited growth conditions (9,11,12,29,30). It has been suggested that the common factor leading to increased GDGT cyclization at increased temperature, decreased pH, and increased electron donor or acceptor limitation is chronic energy stress on cells (10,29).

The relationship between environmental conditions, namely pH and temperature, and lipid cyclization informs interpretations of GDGT biomarkers in modern and ancient sedimentary records (21,22,31–33). Correlation between the degree of cyclization (relative abundance of rings) and sea surface temperature was demonstrated in marine sediments and forms the basis for the paleotemperature proxy TEX86 (32). However, this relationship is not observed across all environments, particularly in thermal springs (34–37) and at low and high latitudes in marine sediment calibrations (21,38–40). Furthermore, GDGTs from an ecotype of deeply-sea ammonia-oxidizing Archaea (AOA) do not follow the same temperature-dependent trend as those from shallower water AOAs (41). It has also been shown that a combination of pH and temperature correlated with the degree of GDGT cyclization in samples from a wide array of thermal springs (35–37).

The enzymes responsible for introducing cyclopentyl rings into GDGT lipids were only recently discovered. Zeng and colleagues identified the first known GDGT ring synthase genes, *grsA* and *grsB*. These genes code for enzymes GrsA and GrsB, which catalyze cyclopentyl ring formation in isoprenoid lipid precursors in two model organisms, *Sulfolobus acidocaldarius* and *Saccharolobus solfataricus* (42). Specifically, GrsA and GrsB catalyze the formation of between one and four rings each, for a total of up to eight rings. Each enzyme acts at different locations on the hydrocarbon chain. GrsA catalyzes the introduction of the first suite of rings for GDGT-1 to -4, whereas GrsB cyclizes ring introduction at locations where a ring was already formed by GrsA, enabling the formation of GDGT-5 to -8. Because *grsA* and *grsB* share ancestry, we refer to all related homologs as part of a “*grs”* gene family.

To better understand the distribution of GDGT cyclization capacity across environments representing multiple extremes, we surveyed *grs* genes in microbial populations from terrestrial hot spring (meta)genomes and isolates. Hot springs exhibit wide gradients in pH and temperature, two key parameters demonstrated to have a strong influence on GDGT cyclization in pure culture experiments and that show strong correlations to bulk community GDGT profiles in environmental samples. As such, we predict that these parameters have substantially influenced the diversification of *grs* in hot spring communities. Hot springs are home to most known major lineages of the Archaea (43,44), which enables a robust survey of *grs* genes across taxonomic groups. In this work, we address the following question: Is the distribution and abundance of *grs* genes correlated with spring pH, temperature, or other geochemical parameters? To do so, we queried isolate genomes, single cell genomes, metagenomes, and transcriptomes from a global compilation of terrestrial hot springs for *grs* homologs. These genes were subjected to a variety of informatics analyses to identify patterns in their distribution as a function of environmental parameters. Further, the *grs* genes were subjected to phylogenetic analyses to examine the role of pH, temperature, and geochemistry in shaping their evolutionary history. Our findings provide insights into the importance of environmental pressures on microbial membrane cyclization potential in extreme environments and the role cyclic GDGTs have played in archaeal diversification across the Earth systems.

## 2.0 Methods

### 2.1. Reference sequence database curation

We compiled a reference dataset of 35 amino acid sequences of *grsA* and *grsB* genes functionally characterized in *Sulfolobus acidocaldarius* and *Saccharolobus solfataricus* (42) and of their homologs in other cyclic GDGT-producing Archaea (File S1) using the following procedure. Cyclic GDGT-producing Archaea were identified based on previously published lipid profiles (22,28). At least one genome from each order-level lineage of known cyclic GDGT-producing Archaea was selected. Homologs were then detected by searching genomes of these Archaea with *S. acidocaldarius grsA* and *grsB* sequences as queries using BLASTp (45) and requiring ≥70% coverage, ≥20% identity, and E-value ≤10^−40^ to both query sequences, which are similar cutoffs to those used by Zeng and colleagues (42). In cases where a genome possessed more than one sequence matching these criteria, all candidate sequences were included in the reference set. Reference sequences and BLASTp results are reported in Table S1.

### 2.2. Compilation of datasets

In total, 1,396 thermal springs datasets, consisting of archaeal and bacterial isolate genomes, metagenomes, single cell genomes, metatranscriptomes, and metagenome-assembled genomes (MAGs) that contained pH and/or temperature metadata, were selected from the Joint Genome Institute’s (JGI) Integrated Microbial Genomes and Microbiomes (IMG/M) resource (last access July 2021, Table S2 and S3)(46). From within these datasets, 1,114 high quality MAGs (≥80% estimated completeness and ≤5% estimated contamination) derived from 61 of the metagenomes were available for analysis. Specifically, 799 MAGs are from 34 YNP metagenomes, while 315 MAGs, identified by the automated binning pipeline in IMG, are from 27 metagenomes (Table S9, sources reference therein). MAG quality was predicted based on CheckM estimates (47). The location of these datasets is outlined in the Data Availability statement below.

### 2.3. Identification of *grs* homologs

The thirty-five sequences in our reference set were aligned in MAFFT (v. 7.480) using the L-INS-i model with default parameters (48). A Hidden Markov Model (HMM) profile was built from the alignment using the hmmbuild function in HMMER v. 3.1b2 (49). The HMM profile was used to query the assembled and inferred protein files for the 1,396 hot spring datasets, using hmmsearch. The HMM search was automated using Snakemake v. 6.4.1 as a workflow engine (50). Matches with an e-value ≤10^−40^ and a length of at least 400 amino acids were retained. The stringent length threshold was used to ensure conservative designation of *grs* homologs. Sequences meeting this threshold criterion aligned well with the reference sequences across multiple domains. The HMM profile was augmented using a bait and refinement approach, including 25 homologs detected in genomes of isolates from thermal springs during the first HMM search (File S2) (51). All homologs detected using the refined profile and meeting threshold criteria were used in downstream analyses. Contig identifiers were then used to associate *grs* sequences to the selected MAGs.

### 2.4. Abundance analysis

The relative abundance of *grs* copies within a metagenome was estimated using recovered *grs* sequences, normalized for both coverage and read depth for metagenomes with reported coverage data (n = 64 metagenomes). The coverage of each *grs* gene was estimated using average coverage of the associated scaffold (in reads). These coverage values were summed for all *grs* sequences in a metagenome to get the total coverage and divided by the total mapped reads per million. Relative abundance was reported in reads per million (RPM).

A multiple regression analysis was used to evaluate the relationship between, pH, temperature, and *grs* abundance. Multiple regression models were tested and evaluated based on the Akaike Information Criterion (AIC) score, which is used to compare the likelihood of different statistical models, and adjusted-R^2^ values, which describe the proportion of the variance explained by the model. For a subset of metagenomes from Yellowstone National Park (YNP) (n = 34), additional geochemical parameters (dissolved oxygen, ferrous iron, sulfide, sulfate, chloride) were available and were included and tested for Pearson’s correlation with pH and/or significance in linear regression models. Regression analysis was performed in *R* (v. 1.4.1717) using the packages *car* (R > v. 3.0), *mosaic* (v. 1.8.3), *tidyverse* (v. 1.3.1), and MuMIN (v. 1.43.17).

### 2.5. Copy Number

Copy numbers of *grs* for MAGs and isolates were calculated by summing the number of putative *grs* homologs recovered during HMM searches. In cases of individual MAGs, copy number may be underestimated due to incomplete MAGs and the use of a stringent length filter for *grs* identity.

Kruskal-Wallis non-parametric one-way analysis of variance with ranks was used to determine if there was a significant difference between temperature and pH distributions with respect to isolate copy number or *grs* presence and absence. Growth conditions for isolates were extracted either from the associated publication on the Genomes OnLine Database (52), or, if not available there, from the publications describing the type strains in the *International Journal of Systematic and Evolutionary Microbiology* (IJSEM) or BacDive (53) (Table S8). Dunn’s test was performed *post hoc* to determine which pairwise comparisons were significant, using Bonferroni P-value adjustments and an alpha significance threshold of 0.05. The same analysis was performed on MAGs, using pH and temperature of the associated samples. Statistical tests were performed with the SciPy statistics package v. 1.9.0 (54).

### 2.6. Phylogenetic analysis

From the 2,815 putative *grs* amino acid sequences identified in the HMM search across all thermal springs data sets, 2,112 were unique (i.e., defined at a 100% amino acid sequence identity threshold) and were used for phylogenetic analyses. A multiple sequence alignment was created for the unique sequences in MAFFT (v. 7.480) using a fast Fourier transform method with iterative refinement (FFT-NS-i) (48). The phylogenetic tree was reconstructed in RAxML (v. 8.2.12), specifying LG+G for the amino acid model with empirical base frequencies and gamma model, respectively, for rate heterogeneity with 100 bootstrap pseudosamples (55). The outgroup (n = 36 sequences) was formed with homologs to an *S. acidocaldarius* radical SAM protein (tRNA uridine(34) 5-carboxymethylaminomethyl modification radical SAM/GNAT enzyme Elp3; GenBank AAY80401.1) with distant homology to GrsA and GrsB. Additional homologs for the outgroup were identified by a BLASTp search of the *S. acidocaldarius* radical SAM protein against the NCBI *nr* database, and a subset of sequences was selected to cover a range of archaeal groups (coverage ≥80%, E-value ≤10^−60^). This outgroup was selected after testing several radical SAM proteins for the one producing the most consistent tree topology and highest bootstrap support. The tree was annotated with the pH and temperature values of the sample associated with each *grs* homolog (52). Correlation between the evolutionary history of *grs* homologs and distribution of pH and temperature values was evaluated using the Pagel’s lambda (λ) metric (56) as implemented in the *Phylosig* (Phytools package version 0.7-90) package in R (57). A likelihood ratio test implemented in *Phylosig* was used to assess significance of the deviation from the null model (lambda = 0).

A second phylogeny was constructed using a subset of *grs* protein sequences associated only within the selected MAGs in order to relate *grs* sequences to taxonomic affiliation and copy numbers within MAGs (47). To reduce the size of the dataset, sequences were clustered at 99% amino acid identity using cd-hit v. 4.6.1 (58), with “-g 1 -c 99”options, and the longest sequence was kept as the representative for each cluster, leaving 315 sequences. *Grs* protein sequences from thermal spring isolates (n = 53) and other cultured Archaea (n = 24) were also included for reference. The alignment and phylogeny were constructed with the same method as described above.

Both trees were visualized in iTOL v. 6.5.2 (59). Each leaf was annotated with the taxonomic class predicted by the Genome Taxonomy Database Toolkit (GTDB-tk, v. 1.3.0) (60) and the number of *grs* protein sequences detected in the associated MAG.

A naïve Bayes classifier was developed to categorize sequences into GrsA-like and non-GrsA-like groups using their physicochemical properties (inspired by Kogay and colleagues (61)). Physicochemical properties of the amino acids in each sequence were tallied and summed (Table S12). The model was trained on a set of protein sequences closely related to GrsA and a set of protein sequences closely related to GrsB. The model was used to classify all other sequences either as GrsA or GrsB. The classification was overlaid on the tree of Grs homologs from MAGs (Figure S1, Table 13).

## 3.0 Results

### 3.1. Homologs of *grs* are present in geothermally-influenced springs across a wide range of temperature and pH

Among 1,396 thermal spring datasets, 2,112 unique *grs* homologs were detected in 307 datasets comprising 243 metagenomes, 39 isolate genomes, 11 single cell genomes, and 14 metatranscriptomes (Table S3). These 307 datasets derive from seven countries spanning four continents, listed in order of most to least samples: U.S., China, Canada, New Zealand, Russia, Japan, and Portugal (Table S4). Samples from the U.S. are from springs in Wyoming, California, Nevada, and Oklahoma, with the majority from YNP, where *grs* homologs were detected in 132 samples out of 183 YNP samples queried. The geothermally influenced springs come from a range of geologic settings, including volcanic hotspots, regions of tectonic plate subduction and collision, as well as in regions impacted by significant faulting (64–67). Across these 307 datasets, 2,815 *grs* homologs were detected in sites that spanned a temperature range of 10 to 98 °C (median 77 °C) and pH of 1.30 to 9.60 (median 6.37) (Figure 1, Table S5).

**Figure 1.**
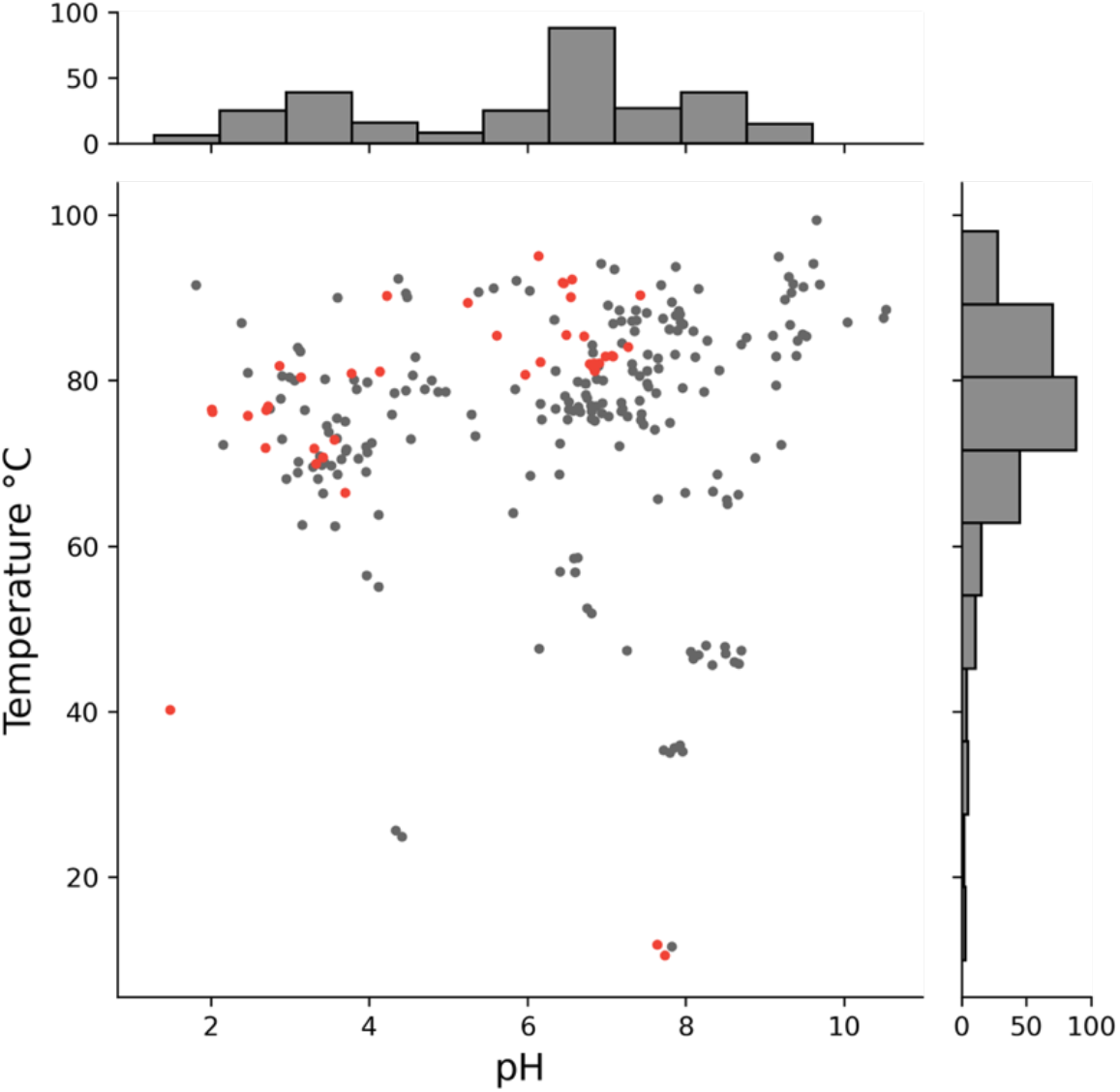
Distribution of temperature and pH among the 307 thermal spring datasets in which *grs* homologs were detected. Thermal spring isolate and single cell genomes are highlighted in red. Marginal histograms show the number of datasets.

### 3.2 pH and temperature predict *grs* distribution and abundance

The relative abundance of *grs* was assessed in all metagenomes with coverage data (n = 64). In thermal springs across the world, pH and temperature are significantly correlated with the relative abundance of *grs*. Specifically, around 60% of the variance in the relative abundance of *grs* is explained by spring pH alone (adjusted R^2^ = 0.602, *P* = 2.98 × 10^−14^, df = 62, AICc = 323.6), where lower pH is associated with higher *grs* abundance (Figure 2). Adding temperature to the model led to predictive power capturing 65% of the variation in *grs* abundance (adjusted R^2^ = 0.647, *P* = 6.29 × 10^−15^, df = 61, AICc 321.8). Higher temperature springs are significantly associated with higher *grs* abundance, though the correlation is not as strong as for pH. Alone, temperature explains just over 10% of the variance in *grs* abundance (adjR^2^ = 0.106, *P* = 0.005). For every unit decrease in pH, the relative abundance of *grs* increases by 15.44 RPM (95% CI: 12.29 to 18.61 RPM) and for every 10 °C increase in temperature, *grs* abundance increases by 5.78 RPM (95% CI: 1.87 to 9.69 RPM) (Figure 2). The difference in the sum of squares for each variable shows that pH accounts for around 12 times more variation in *grs* abundance than temperature does (Sum of squares: pH = 61,589, Temperature = 4,957).

**Figure 2.**
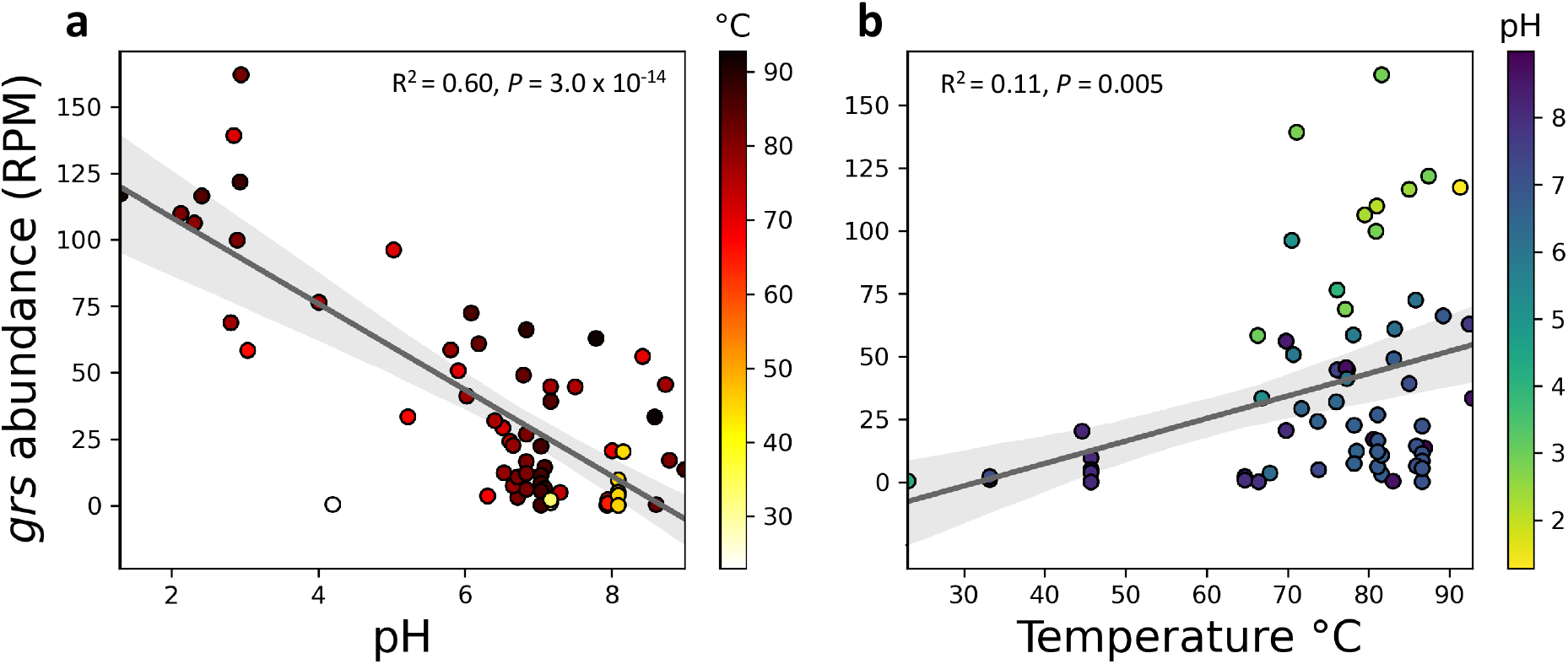
Correlations between *grs* homolog relative abundance and sample site pH (a) or temperature (b) in 64 hot springs metagenomes. The relative abundance of grs is reported in reads per million (RPM). Temperature and pH are significantly correlated with *grs* relative abundance (multiple linear regression model, adjusted R^2^ = 0.647, *P* = 6.29 × 10^−15^, df = 61, AICc = 321.8). Individual linear regressions of pH and temperature are indicated on the plots and shaded regions represent 95% confidence intervals.

We tested for correlations between *grs* homolog abundance and concentrations of dissolved oxygen, ferrous iron, chloride, sulfate, and sulfide in a subset of YNP hot springs for which such data was available (n = 34 metagenomes). Oxygen, iron, and sulfide were not found to have significant associations with *grs* abundance. Springs with higher sulfate concentrations, or higher sulfate to chloride ratios, correlate strongly with lower pH, consistent with prior observations (68). As a result, sulfate concentrations and sulfate:chloride ratios also positively correlate with *grs* homolog abundance (Figure S2).

### 3.3 Genomes from the most acidic environments contain multiple copies of *grs*

While the relative abundance of *grs*, measured in RPM, describes its prominence within a microbial community, the *grs* homolog copy number in a genome or MAG describes the abundance of the gene at the organismal scale. In archaeal isolates, there was a significant difference among distributions of optimal growth pH and the number of *grs* homolog copies (e.g., one, two, or more) per genome (Kruskal-Wallis: *P* = 1.2 × 10^−4^, post hoc Dunn’s test: copy numbers 1 and 2: *P* = 1.9 × 10^−4^; 1 and 3: *P* = 0.03) (Figure 3a,b). The genomes of all archaeal thermal spring isolates with a low pH growth optimum (pH < 4) encode more than one copy of *grs*. The pH distributions of archaeal MAGs with *grs* copy numbers of one and two are significantly different (Kruskal-Wallis, *P* = 2.2 × 10^−20^; post hoc Dunn’s test, *P* = 9.7 × 10^−7^) (Figure 3c). MAGs with two copies of *grs* are more widely distributed at lower pH than those with just one copy.

**Figure 3.**
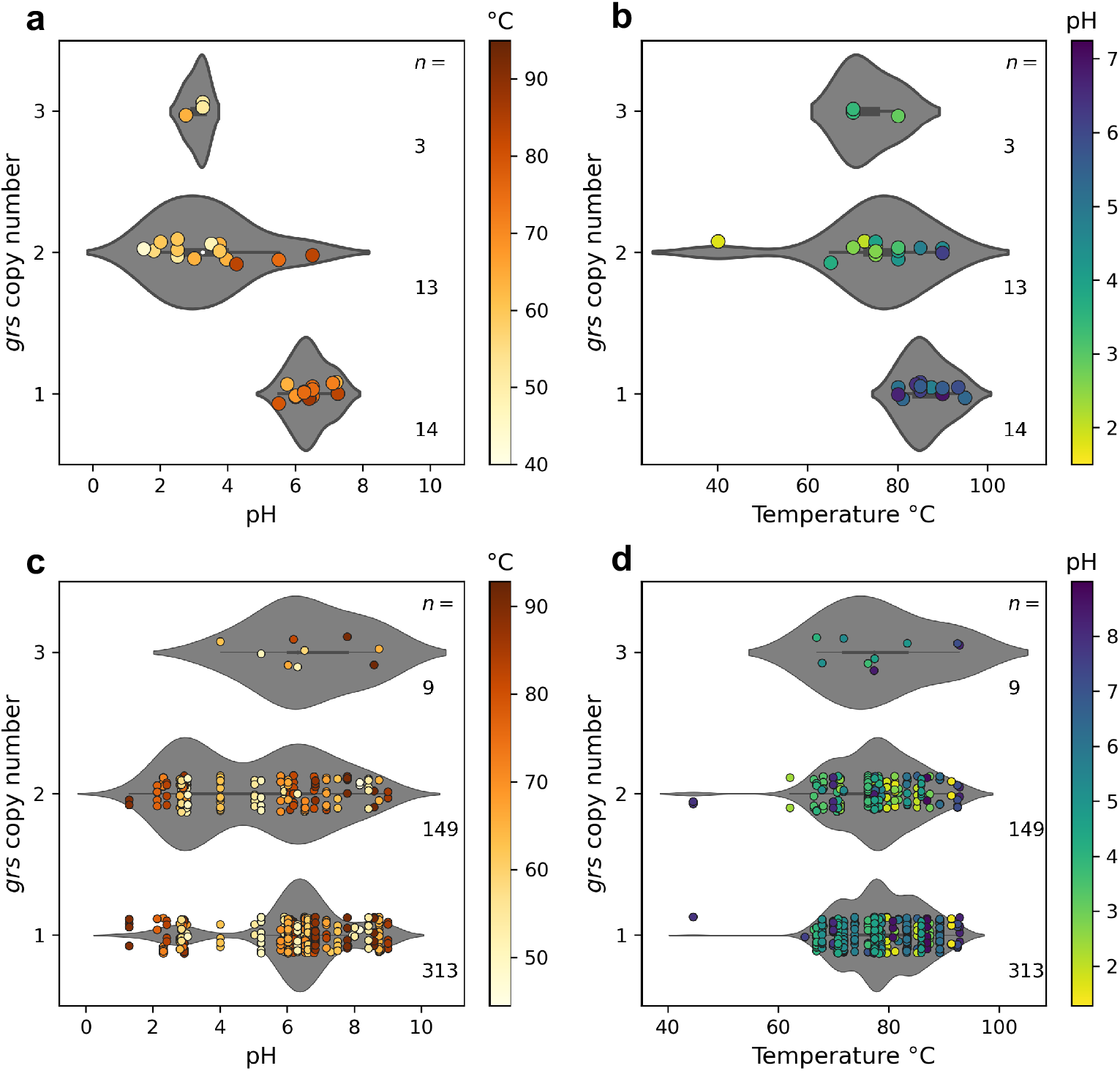
Copy numbers of *grs* in isolates and MAGs across hot springs of varying pH and temperature. The optimal growth pH (**a**) and temperature (**b**) versus *grs* copy number across 30 thermal spring archaeal isolates. Midpoints are used when optimal growth pH or temperature were reported as a range (pH: n = 15 isolates, temp: n = 8 isolates), otherwise, individual values are used. The environmental pH (**c**) and temperature (**d**) versus *grs* copy number across 474 high quality *grs-*encoding MAGs. MAGs with copy numbers of five (n = 2, pH = 7.2, 7.5, Temperature = 76.1, 77.4) and six (n = 1, pH = 7.3, Temperature = 73.8) are not shown. Violin plots represent the distribution using the kernel density estimate and are normalized to the same width.

The temperature distributions of archaeal isolates whose genomes encode one versus two *grs* copies are significantly different (Kruskal-Wallis: *P* = 3.6 × 10^−3^, post hoc Dunn’s test: *P* = 4.0 × 10^−3^). However, many of the isolates with two copies of *grs* are from more acidic environments than those with one copy, including *Acidiplasma aeolicum* (V), which has a copy number of two and grows at very low pH (1.5) and in mesophilic environments (40 °C) (Table S8). This makes it difficult to assess the relationship between temperature and copy number independent of pH. The genomes of archaeal thermal springs isolates that have high temperature growth optima (T > 70°C) encode anywhere from one to three copies of *grs*. The temperature distributions across MAGs of the various copy numbers are not significantly different (Figure 3d)

### 3.4 *grs* copy number per genome varies across taxonomic classes

Among the 1,114 high-quality MAGs, 581 were archaeal and 533 were bacterial (Table S9). We detected one or more copies of *grs* across 501 MAGs (n = 497 archaeal and n = 4 bacterial) comprising a total of 690 sequences (Table S10). The *grs*-encoding MAGs derive from 12 out of 47 archaeal classes and one bacterial class (Acidobacteriae) (Table S7, Table S10). The archaeal class Thermoproteia (n = 424 sequences from n = 297 MAGs) is the most well represented. Within the Thermoproteia, those associated with the order Desulfurococcales (n = 156 MAGs) are the most abundant among *grs*-encoding MAGs. We also detected homologs of *grs* in several archaeal groups which lack some or any cultured representatives, including the classes Korarchaeia (prev. phylum Korarchaeota; n=12 MAGs), Bathyarchaeia (prev. Phylum Bathyarchaeota; n = 30 MAGs), and Hadarchaeia (prev. Hadesarchaeota; n = 11). Homologs of *grs* were also found in several thermal spring isolate genomes for which lipid composition has not been studied, to our knowledge. These include members of the genus *Thermofilum* (order Thermofilales) and *Pyrobaculum* (order Thermoproteales) (Table S8).

The distribution of spring pH from which archaeal *grs*-encoding MAGs were detected is significantly different from the distribution of spring pH where MAGs lacked detectable *grs* homologs, with *grs-*encoding MAGs more widely distributed at low pH (Kruskal-Wallis, *P* = 0.0019) (Figure S3). Conversely, the distribution of spring temperatures, from which archaeal *grs*-containing MAGs were found, is not significantly different from spring temperatures for which *grs* was not detected (*P* = 0.40).

Most MAGs encode one (n = 336) or two copies of *grs* (n = 151), and only 14 MAGs encode three or more copies (Figure 4). The *grs* copy number varies both between and within taxonomic groups at the class and order level (Figure 4 and S4). Three copies were detected amongst members of the classes Thermoproteia (n = 6 MAGs), Nitrososphaeria (n = 2), and Hadarchaeia (n = 1). Four copies (n = 1 MAG), five copies (n = 3), and six copies (n = 1) are present in representatives from the order Nitrososphaerales (c. Nitrososphaeria).

**Figure 4.**
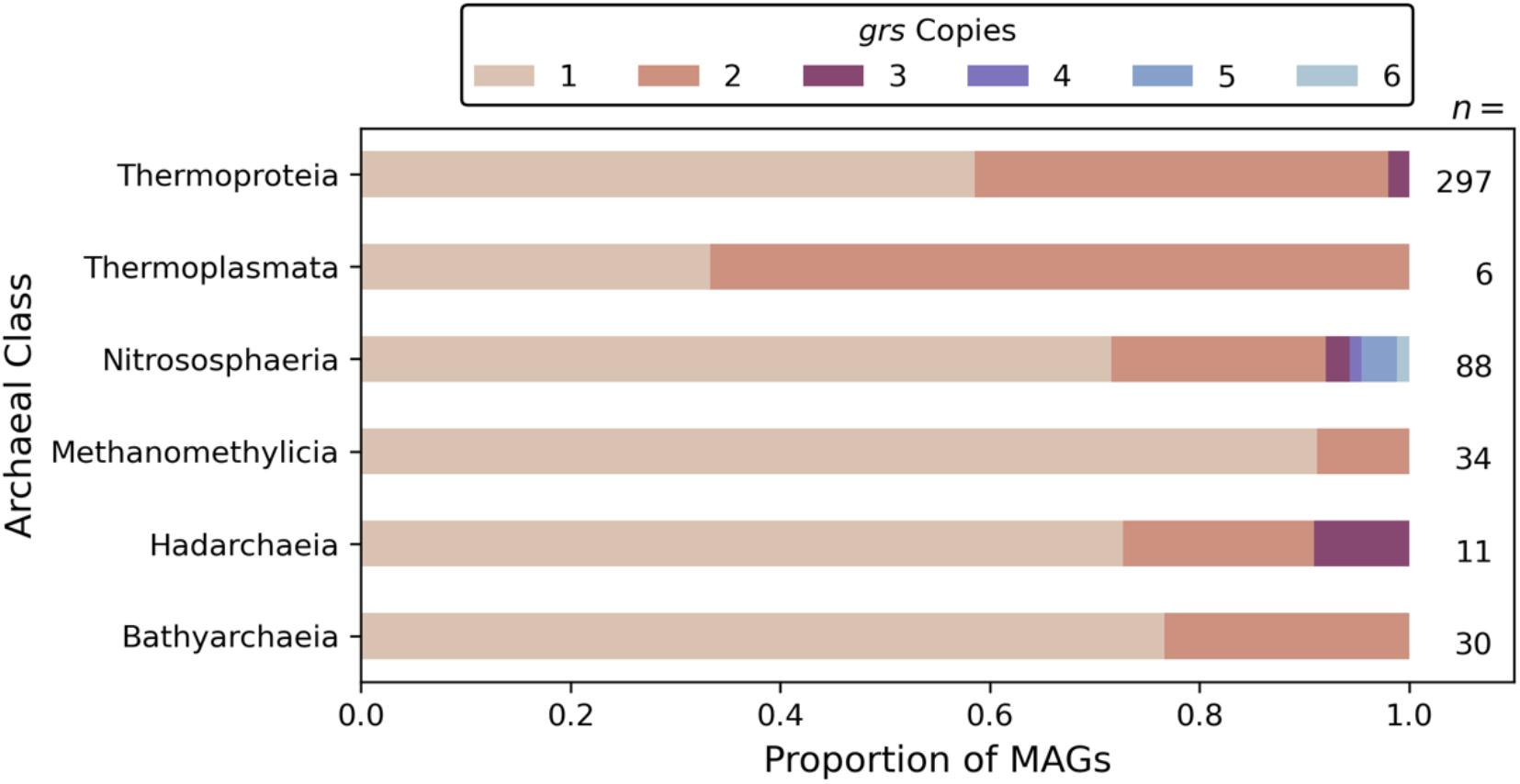
Variation in *grs* copy number across taxonomic classes. The copy number of *grs* varies with taxonomic class in 501 MAGs assembled from thermal spring metagenomes. Classes with only one copy detected across all surveyed members are not shown: archaeal classes Micrarchaeia (n = 1), Methanocellia (n = 1), Lokiarchaeia (n = 1), Korarchaeia (n = 12), Archaeoglobi (n = 16); and bacterial class Acidobacteriae (n = 4).

In the framework established by Zeng and colleagues (2019), the *S. acidocaldarius* genome encodes *grsA* and *grsB* (*grs* copy number of two), whereas the *S. solfataricus* genome encodes one *grsA* and two *grsB* homologs (*grs* copy number of three). Importantly, Zeng and colleagues also found that the ring cyclase GrsB preferentially acts on the substrates produced by GrsA, although it can also produce rings on the acyclic GDGT-0 substrate, albeit in smaller quantities.

### 3.5 Phylogenetic analysis suggests a complex evolutionary history of the *grsAB* gene family

The phylogenetic relationships among the 2,112 Grs protein sequences show that there is a substantial diversity beyond the experimentally characterized homologs (Figure S5). Overall, homologs do not universally cluster by pH or temperature, although some small clusters by pH are apparent. To test if these clustering patterns are only a consequence of the evolutionary history of the *grs* gene homologs, we transformed the tree using Pagel’s λ (56) and pH and temperature as traits and assed the phylogenetic correlation to either parameter. For both pH and temperature, the null model of the independent evolution of these traits from phylogeny was rejected (pH: λ = 0.88, *P* = 0; temperature: λ = 0.93, *P* = 1.09 × 10^−266^), suggesting an association of both pH and temperature with phylogenetic position of Grs.

A smaller phylogeny of Grs homologs found only in MAGs reveals a complex evolutionary history of this gene family (Figure 5). The tree does not have a topology that follows established taxonomy, suggesting horizontal gene transfer events (69). Homologs from the same MAG do not usually group together. This suggests that the presence of multiple *grs* homologs in a given genome is not due to recent gene duplication. This is exemplified by the many MAGs from the class Thermoproteia that have two or three copies of *grs* (n = 114); homologs from the same MAG fall on different regions of the tree, suggesting two distinct functional subtypes (i.e. GrsA and GrsB).

**Figure 5.**
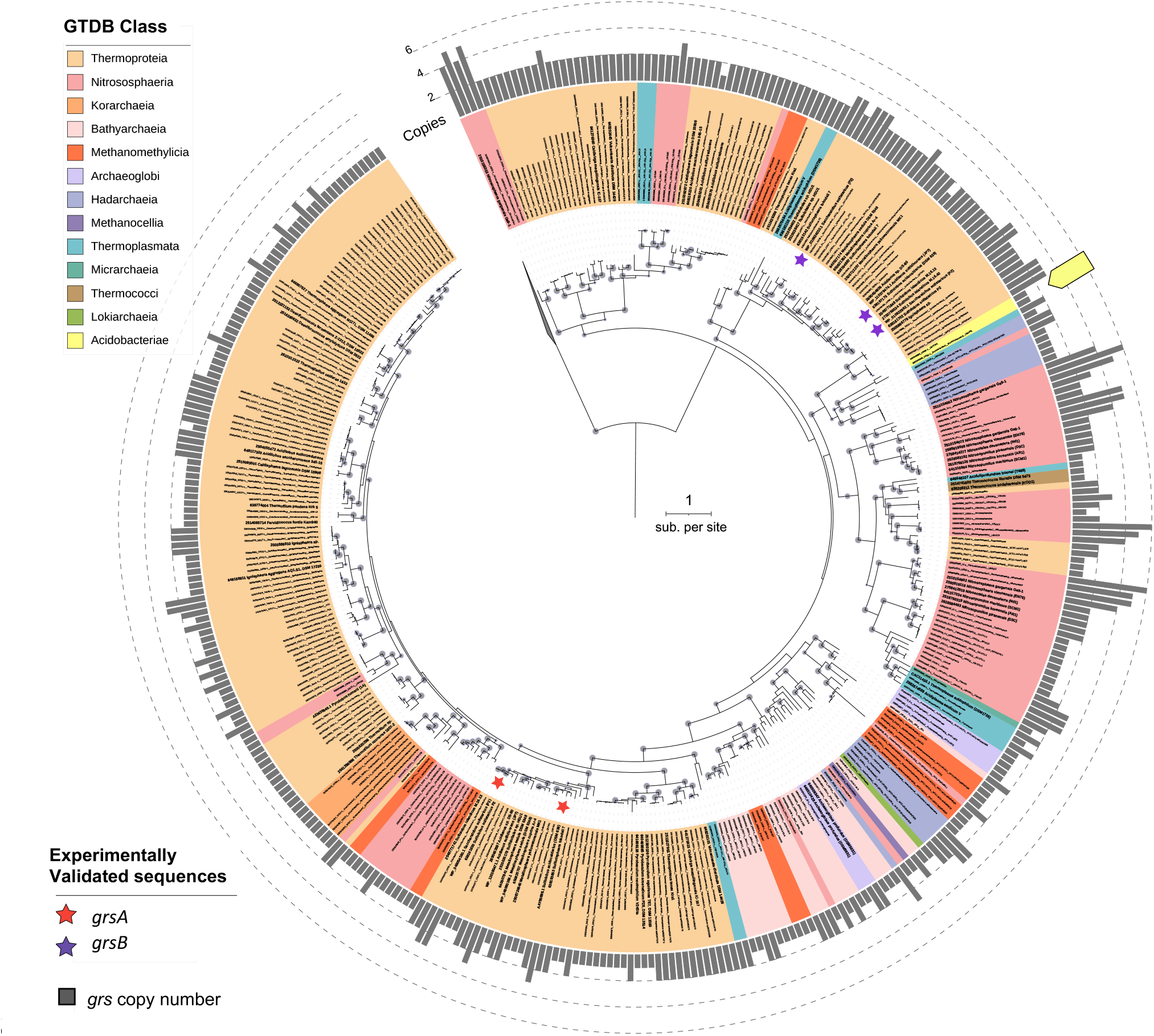
Phylogeny of *grs* homologs from thermal spring MAGs. Rooted maximum likelihood phylogeny of 315 putative *grs* protein sequences from 256 MAGs. Experimentally validated *grsA* and *grsB* protein sequences (42) are marked by stars. Sequences from thermal spring isolates (n = 53) and other cultured Archaea (n = 24) are included as references and are shown in bold. The *grs* copy numbers of the MAGs or genomes associated with each of the sequences are displayed as bars around the sequence terminals. Acidobacteriae sequences are marked by a yellow flag. Bootstrap values ≥ 75 % are marked by gray circles. Tree scale is in units of substitutions per site.

Experimentally validated GrsA and GrsB homologs are distantly related and appear in two separate clades of the tree (Figures 5 and S4). Interestingly, most taxa that group in the clade with the experimentally validated GrsB homologs from *S. sulfolobus* and *S. solfataricus* have two or three *grs* homologs per genome, while many taxa that group in the clade with experimentally validated GrsA homologs have only one *grs* homolog (Figure 5). A machine learning classifier based on the physicochemical properties of amino acid sequences did not distinctively divide GrsA-like and GrsB-like sequences into two separate clades (Figure S1). However, a clade of close homologs to GrsB do share a conserved coiled-coil domain at the N-terminus, which was not detected among other sequences in the phylogeny (Figure S1). Given that the coiled-coil domain is conserved across all sequences in this clade, it may be a useful diagnostic criteria for identifying *grsB* homologs in genomes of thermoacidophiles.

### 3.6 Thermophilic Nitrososphaeria encode up to six *grs* homologs

The class Nitrososphaeria encompasses both terrestrial and marine groups, many of whom are ammonia-oxidizers. Most Nitrososphaeria MAGs encode one or two copies of *grs*. However, exceptionally high *grs* copy numbers (i.e., four to six homologs) were detected in several Nitrososphaeria MAGs (n = 5) (Figure 4). The MAGs with high *grs* copy number come from thermal springs in Yellowstone National Park and northern Nevada in the U.S., and Tengchong in China, spanning 70 to 80 °C, and pH 6.4 to 7.5 (Table S9). Many of these *grs* homologs are closely related and appear in one clade on the tree (Figure 5).

### 3.7 Acidobacteriae MAGs encode *grs* genes

One copy of *grs* was found in each of four MAGs from members of the Acidobacteriae – a class of Bacteria. These MAGs are from moderately acidic (i.e., pH = 5) and circumneutral (i.e. pH = 8) springs in Yellowstone National Park, U.S., and Dewar Creek, British Columbia, and all are predicted to be ≥95% complete with 0% contamination (Table S9). These Grs homologs have significant sequence similarity to the *S. acidocaldarius* GrsAB sequences (BLASTp: *e*-value ≤10^−40^; ≥87 % coverage to GrsA with ≥27% identity; ≥78% coverage to GrsB with ≥30% identity). The bacterial Grs protein homologs are nested within a clade of archaeal homologs, suggesting a possible horizontal gene transfer event from Archaea to Bacteria (Figure 5).

## 4. DISCUSSION

### 4.1 pH is the primary driver of *grs* abundance and distribution

In this study we quantified the distribution and diversity of *grs* homologs from geothermal springs across the planet, focusing on sites from Yellowstone National Park. We found that both pH and temperature are significantly associated with *grs* abundance and are better predictors of *grs* abundance when combined than when applied independently. Spring pH is a stronger predictor of *grs* abundance than temperature and is therefore more likely to indicate the potential for lipid cyclization among Archaea in a given spring (Figure 2). Furthermore, Archaea with the capacity to cyclize GDGTs (i.e., encode one or more *grs* homologs) are more widely distributed in low pH systems than those that cannot (i.e., do not encode *grs*) (Figure S3). Given that lipid cyclization has been interpreted as an adaptation to energetic stress (11,12), these observations are consistent with the understanding that Archaea inhabit environments that impose chronic energy stress (e.g., hot and acidic) (2,70).

Prior culture studies with thermoacidophiles have shown a strong influence of pH on the degree of GDGT cyclization (8,10,71). The supposition that GDGT cyclization enables Archaea to thrive in acidic and hot waters is also consistent with environmental lipid surveys, where several studies document a stronger correlation between hot spring pH and average GDGT cyclization, relative to the relationship between temperature and GDGT cyclization (35–37). In these studies, there was a higher degree of GDGT cyclization observed in acidic springs than neutral or basic springs. The *grs* distribution we describe across wide gradients in temperature and pH strongly supports the conclusion that the capacity for archaeal isoprenoid lipid cyclization was among the key traits that allowed archaeal groups (e.g., the Sulfolobales) to diversify into and dominate acidic and hot environments (36,70). Our gene-level findings for the relationships between *grs* sequences and hot spring or isolate optimal pH and temperature, along with the aforementioned hot spring lipid profiles, consistently point to the primacy of pH in dictating the natural history of *grs*, and ultimately the propensity for GDGT cyclization in a given microbial genome or population.

### 4.2 Multiple copies of *grs* support life in extreme acidity

Environmental pH also plays a role in determining *grs* copy number in a given genome. That is, archaeal MAGs and genomes of isolates with multiple copies of *grs* are more widely distributed in highly acidic environments (Figure 3). We interpret this to indicate that organisms with multiple copies of *grs* are better suited to withstand extreme acidity. For example, *Picrophilus torridus* encodes two copies of *grs* (42) and thrives at a lower temperature than that of the hottest springs (growth optimum ∼ 60 °C), but at a remarkably low optimal pH of 0.7 (72), and produces up to GDGT-8 (10,26). This pattern is reproduced across cultivated thermoacidophiles, where each isolate can generate GDGTs with five or more rings, and each has multiple copies of *grs* (Table S8) (8,10,73,74). Aside from *S. sulfolobus* and *S. solfataricus*, other thermoacidophilic organisms with multiple copies of *grs* include: *Picrophilus oshimae* and the aformentioned *P. torridus* (42), *Acidilobus saccharovorans*, and *Sulfurisphaera tokodaii* (this study, Table S8). This is consistent with the interpretation that GDGTs with higher ring numbers, like GDGT-8, are especially suited to withstand extreme acidity.

Interestingly, isolates from thermal springs who possess only one *grs* copy are not as limited in the range of habitable temperatures, where observations range up to 93.5 °C (Figure 3B), though also live in less acidic sites. Previous work has shown that cyclic GDGTs are not a requirement for life at the highest temperatures in environments that are not acidic (71,75). Further, some hyperthermophilic Archaea have a single copy of *grs*, such as *Methanopyrus* spp., which inhabits circumneutral environments. This aligns with the observation that *Methanopyrus kandleri* can synthesize only up to GDGT-4 when growing near its optima of 100 °C, and can live at temperatures as high 122 °C, but still only at circumneutral pH’s of 6.2 to 6.5 (22,76).

Together these observations suggest acidity has a stronger influence on and is more predictive of *grs* copy number than temperature. Furthermore, we speculate that the evolution of multiple *grs* copies in thermoacidophilic Archaea was an adaptation that allowed ecological expansion into the most acidic and hottest environments on Earth, as has been hypothesized previously (36,70).

### 4.3 Multiple homologs of *grs* with unknown function in the Nitrososphaeria and AOA relatives

A subset of organisms from the class Nitrososphaeria compose the AOA, where all known cultures representatives synthesize a special iGDGT with four cyclopentyl rings and one cyclohexyl ring, known as ‘crenarchaeol’. The enzyme responsible for the 6-memberd ring is still unknown. The GDGTs from these organisms are the basis for a widely applied paleoclimate proxy (32). Genomes and MAGs derived from or closely related to known ammonia-oxidizing Archaea (AOA) commonly have two or more *grs* homologs despite producing only up to four cyclopentyl rings (Figure 4) (28). As such, the function of multiple *grs* homologs in the ammonia-oxidizing Archaea cannot yet be accounted for by our current understanding of Grs enzymatic function, in which one *grs* homolog is responsible for the formation of up to four rings and another for five to eight rings (42). Moreover, all known AOA produce crenarchaeol (28). It is possible that one of the additional *grs* homologs is responsible for the introduction of the cyclohexyl ring in crenarchaeol. To determine if one of these homologs is responsible for crenarchaeol production necessitates a similar approach to that used to characterize *grsAB* in *S. acidocaldarius* and *S. solfataricus* (42).

We observed the highest number of *grs* copies (four to six) among five Nitrososphaerales MAGs from circumneutral environments (Figure 4). Many of these *grs* copies are closely related to one another and may be the result of gene duplication. Recent work with complete genomes shows that large horizontal transfer events with subsequent gene duplications likely drove diversification within the Nitrososphaerales (77). One of the archaeal clusters of orthologous genes annotated at duplication hotspots in the Nitrososphaerales includes *grsA*. Genes for lipid cyclization may have been laterally transferred from other Archaea and/or duplicated as part of a larger pattern of genome expansion during the diversification of the Nitrososphaerales.

The closely related *grs* sequences detected among Nitrososphaerales MAGs and in *Ca*. N. gargensis may be functionally redundant. This is the case for the two *grsB* copies observed in *S. solfataricus*, which perform the same enzymatic function (42). While the two *S. solfataricus* GrsB enzymes both generate GDGT-5 up to GDGT-8, one *grsB* copy was transcribed significantly more than the other. It is likely that environmental or cultivation conditions impact the degree to which particular *grs* sequences are transcribed, though this remains untested.

### 4.4 Horizontal transfer of *grs* from Archaea to Bacteria

While isoprenoid GDGTs are unique to the Archaea, Bacteria are known to produce similar tetraether lipids called branched GDGTs (brGDGTs). In recent work, cultured members of the Acidobacteriae including *Edaphobacter aggregans* and *Candidatus* Solibacter usitatus have been found to make branched GDGTs (brGDGTs) with up to two cyclopentane rings (78–80). A homolog to *S. acidocaldarius* GrsAB was reported and suggested as a candidate for ring formation in *Ca*. Solibacter usitatus (79,80). The microbial producers of brGDGTs in soil have long been unknown, as are the genes involved in ring formation. The *grs* homologs we detected in Acidobacteriae MAGs represent potential lateral transfer events of a ring synthesis enzyme from Archaea to Bacteria and suggest additional *grs*-derived enzymes as a mechanism for lipid cyclization in Bacteria. To confirm that Bacterial homologs perform isoprenoid cyclization requires experimental validation *in vivo* or *in vitro*.

### 4.5 Key drivers of GDGT cyclization vary across environments

In different environments temperature and pH have varying influence on the degree of lipid cyclization. In the ocean, temperature is well-correlated with the degree of lipid cyclization and pH varies minimally (21,22,27,32,81). In hot springs, pH is a much stronger predictor of cyclization (35–37). Beyond pH and temperature, GDGT cyclization can be influenced by factors that also contribute to cellular energy stress, including growth phase, nutrient limitation, oxidative stress, and hydrostatic pressure (9–12,21,29). Lipid packing is advantageous in hot and acidic environments because it enables cells to increase membrane stability and decrease permeability to protons (7,82). For the same reasons, increasing membrane packing via lipid cyclization will enable cells to conserve energy by preventing the loss of reducing power across the membrane. While cyclic-GDGT-producing Archaea living in the oceans do not experience the same extremes of heat or acid stress as their hot springs counterparts, they can experience chronic energy stress in the form of severe electron acceptor or donor limitation, and indeed shifts in GDGT cyclization in AOAs have been documented under such experimental perturbations (9,11). Given the multiple environmental controls on GDGT cyclization with varying degrees of influence, it follows that the *grs* gene family has likely been shaped by distinct combinations of environmental pressures in different geochemical settings. How this plays out in the marine realm, where pH varies significantly less than temperature, remains to be tested, and is a rich target for exploration.

## 5. Conclusions

The observations made here on *grs* distribution and prior observations on lipid composition in hot springs collectively indicate that the evolution of GDGT lipid cyclization through the activity of Grs was a critical step in enabling Archaea to diversify in Earth’s hottest and most acidic environments. The ability to cyclize isoprenoid GDGTs likely first evolved in a thermophile and later diversified to allow for additional cyclization, enabling the organisms to better tolerate multiple environmental extremes, such as high temperature and low pH, and their cumulative influence on the energy state of the cell. Across the hot spring environments we studied here, pH is the most significant driver of *grs* distribution and evolution, where organisms encoding more copies are more widespread at low pH relative to those lacking *grs*. By documenting the diversity of *grs*-encoding MAGs and isolates across hot springs, we further constrain the producers of cyclic GDGT lipids. This work improves our understanding of the key environmental variables that have influenced GDGT production on Earth across space and time.

## 6. Acknowledgements

This work was supported by the National Aeronautics and Space Administration (NASA) New Hampshire Space Grant (NASA-SSW-80NSSC19K0539, to W.D.L. and L.N.B. via Dartmouth College and the University of New Hampshire). L.N.B. was supported by NSF INTERN as an affiliate at the Joint Genome Institute in the Environmental Genomics group (National Science Foundation INTERN Award #1928309, supplemental to NSF-EAR #1928309 (W.D.L). The work conducted by the U.S. Department of Energy Joint Genome Institute (https://ror.org/04xm1d337), a DOE Office of Science User Facility, is supported by the Office of Science of the U.S. Department of Energy operated under Contract No. DE-AC02-05CH11231. YNP metagenomic datasets were generated from support by NASA (80NSSC19M0150 to D.R.C. & E.S.B.). This research was made possible by public datasets stored in the Joint Genome Institute’s (JGI) Integrated Microbial Genomes and Microbiomes (IMG/M) resource, as well as unpublished Yellowstone National Park metagenomes from William Inskeep. We thank Christie Hendrix, Stacey Gunther, and Annie Carlson at YNP for research permits.

## 8. Data Availability Statement

Data for the set of 34 Yellowstone hot springs metagenomes will be permanently available under bioproject accession PRJNA791658. Data (proposals: 10.46936/10.25585/60007560, 10.46936/10.25585/60000876, 10.46936/10.25585/60001042, 10.46936/10.25585/60007294, 10.46936/10.25585/60001099, 10.46936/10.25585/60001182, 10.46936/10.25585/60000772, 10.46936/10.25585/60007475, 10.46936/10.25585/60007482, 10.46936/10.25585/60007520) were produced by the US Department of Energy Joint Genome Institute (https://ror.org/04xm1d337; operated under Contract No. DE-AC02-05CH11231) in collaboration with the user community. A full list of datasets analyzed in this study are publicly available at the JGI’s IMG/M resource and are listed in the Supplementary Material. Code for identification of homologs, machine learning classification, and figure creation is available at https://github.com/lnblum/grs_hotsprings_pub. Figures, *grs* genes, alignments, and tress (in Newick format) were produced in Jupyter notebook using Python packages Seaborn (62) and Matplotlib (63), and archived at FigShare: https://doi.org/10.6084/m9.figshare.c.5874383.v1.

## Supplement Figures

**Figure S1.**
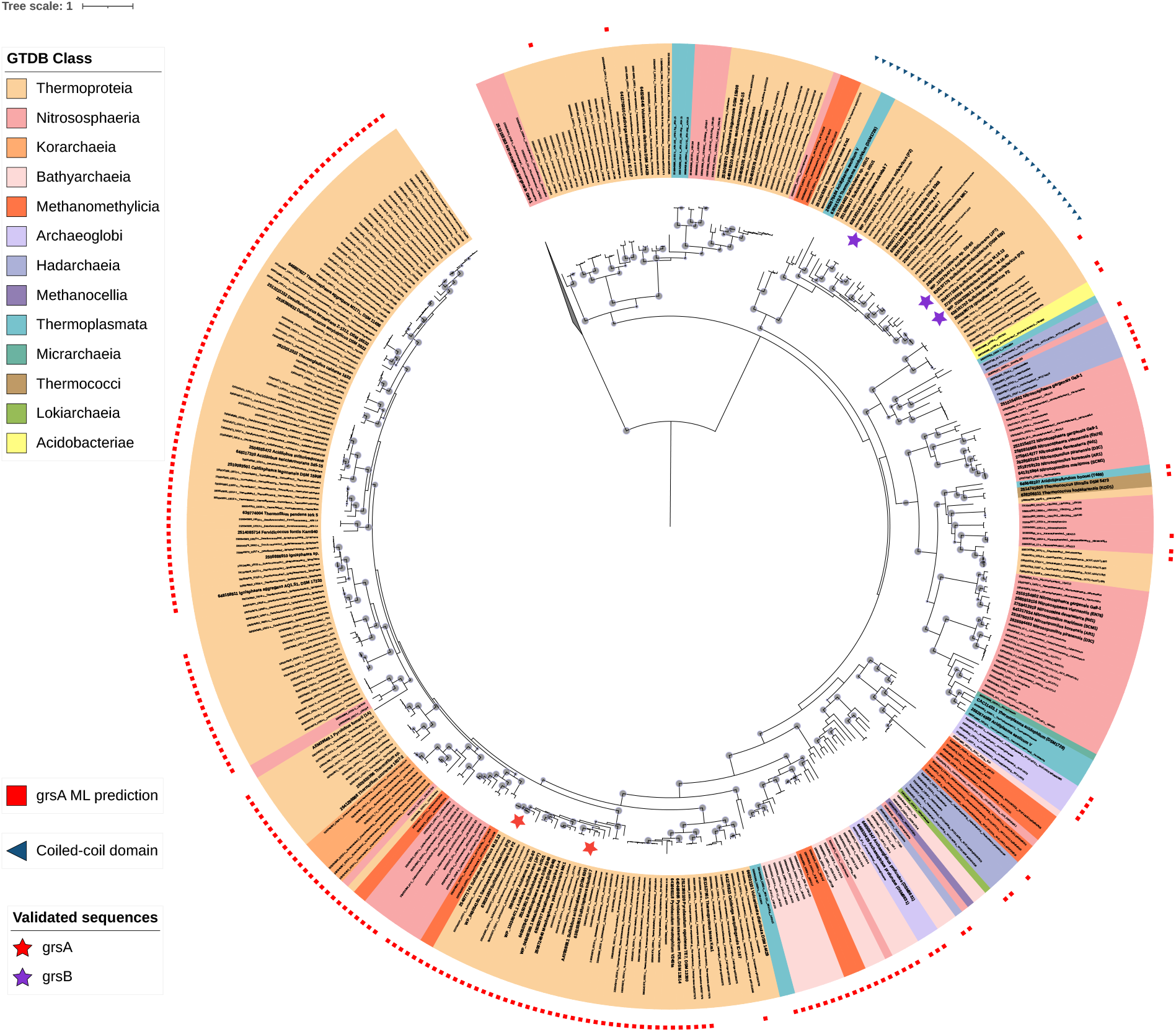
The maximum likelihood phylogeny of *grs* protein sequences depicted in Figure 5, with results of the machine learning classification model predicting *grsA*-like protein sequences overlaid around the outside of the tree.

**Figure S2.**
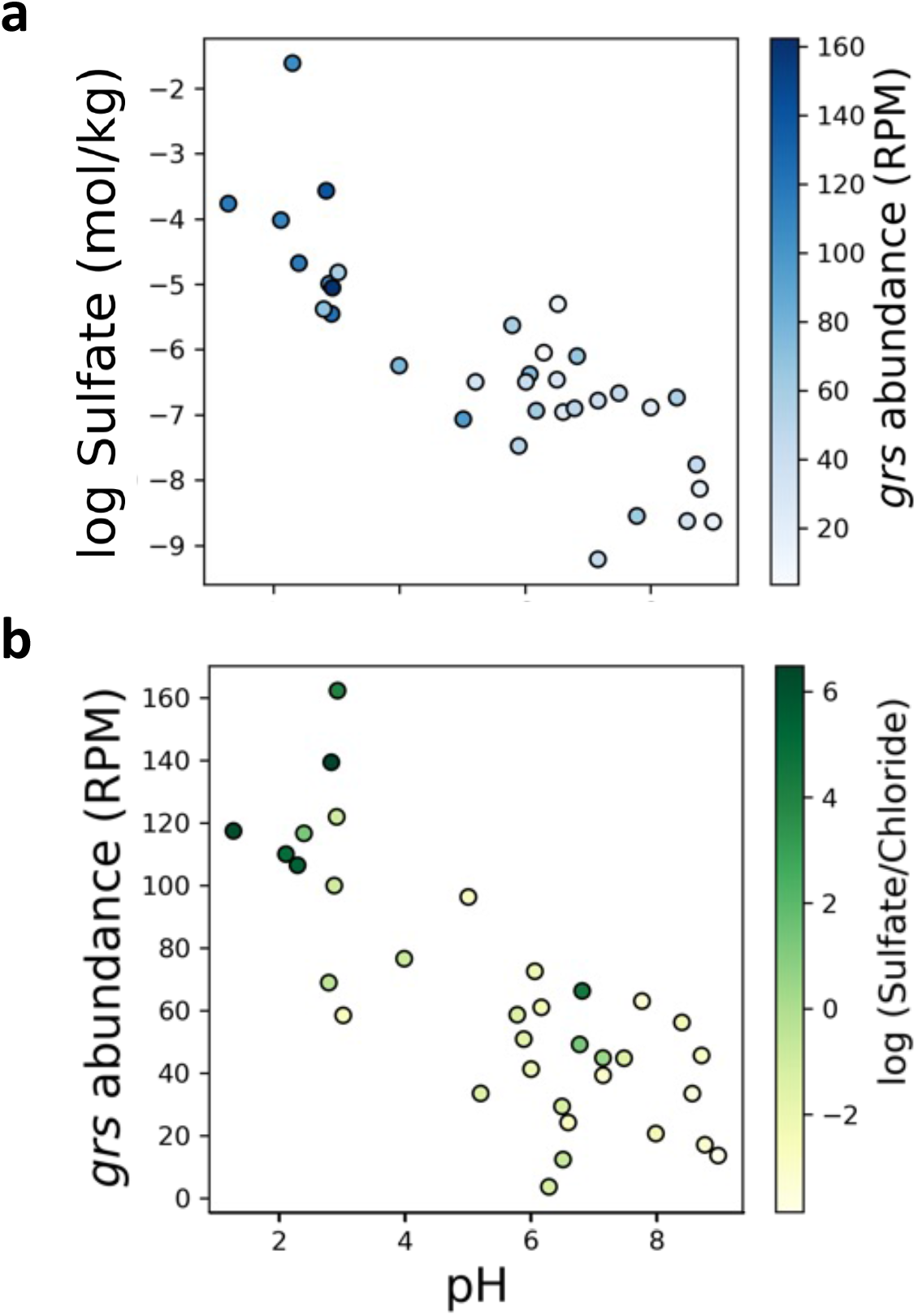
The relationship between select geochemical parameters and *grs* abundance in 34 Yellowstone National Park hot spring metagenomes. A. The negative relationship between pH and log sulfate colored according to *grs* relative abundance. Pearson’s correlation for pH and log sulfate: R = - 0.835 (P < 0.05); *grs* abundance and log sulfate: R = 0.633 (P < 0.05). B. The negative relationship between pH and *grs* abundance, colored according to the sulfate:chloride ratio. Pearson’s correlation for pH and log sulfate:chloride: R = - 0.693 (P < 0.05); *grs* abundance and log sulfate:chloride: R = 0.680 (P < 0.05). Samples for metagenomes were collected in at the same time as geochemical sampling, documented in Table S6.

**Figure S3.**
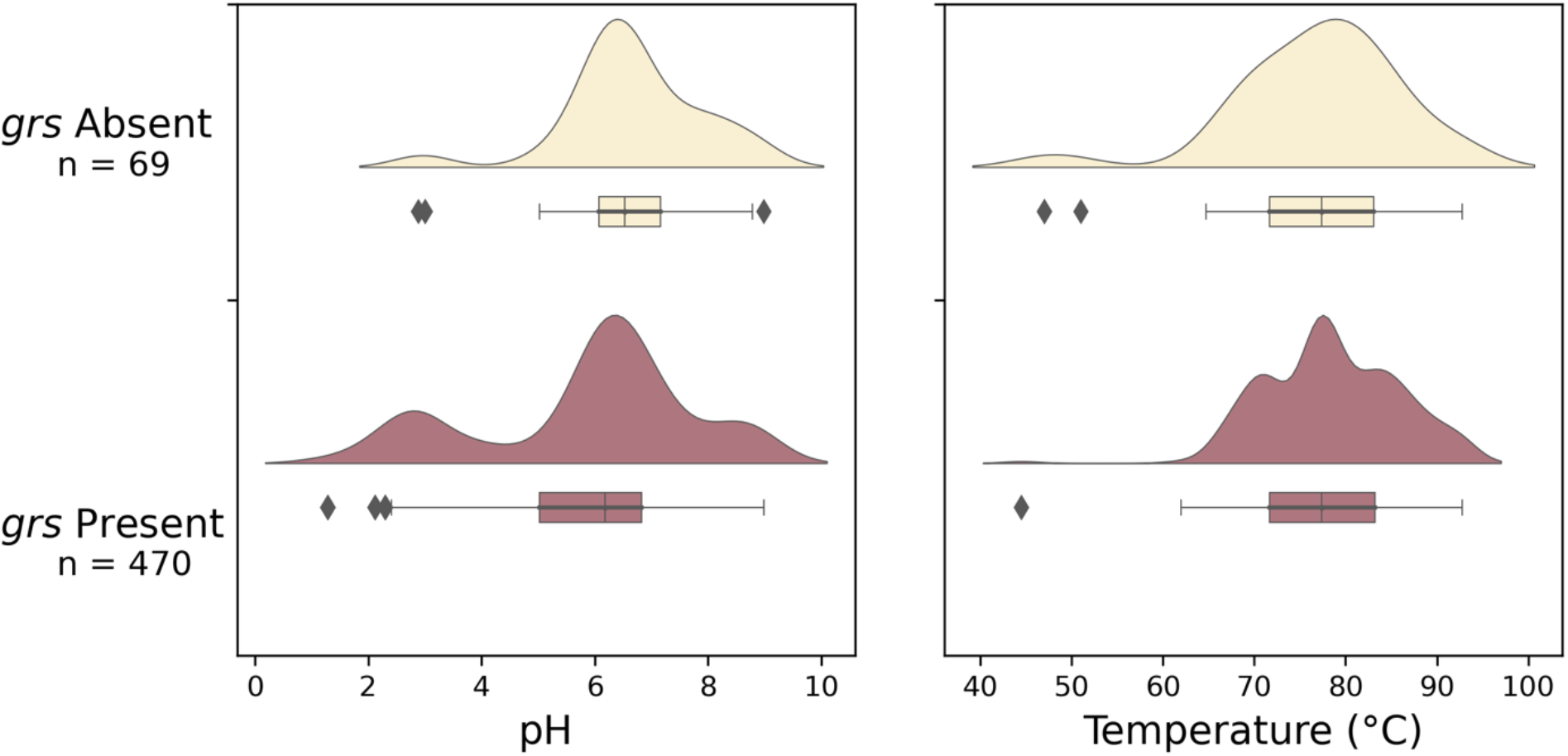
pH and temperature distributions of 539 archaeal MAGs according to whether *grs* is present or absent. Distributions are shown with boxplots and kernel density estimate normalized to the same width. Outliers are defined as data points outside 1.5x the interquartile range.

**Figure S4.**
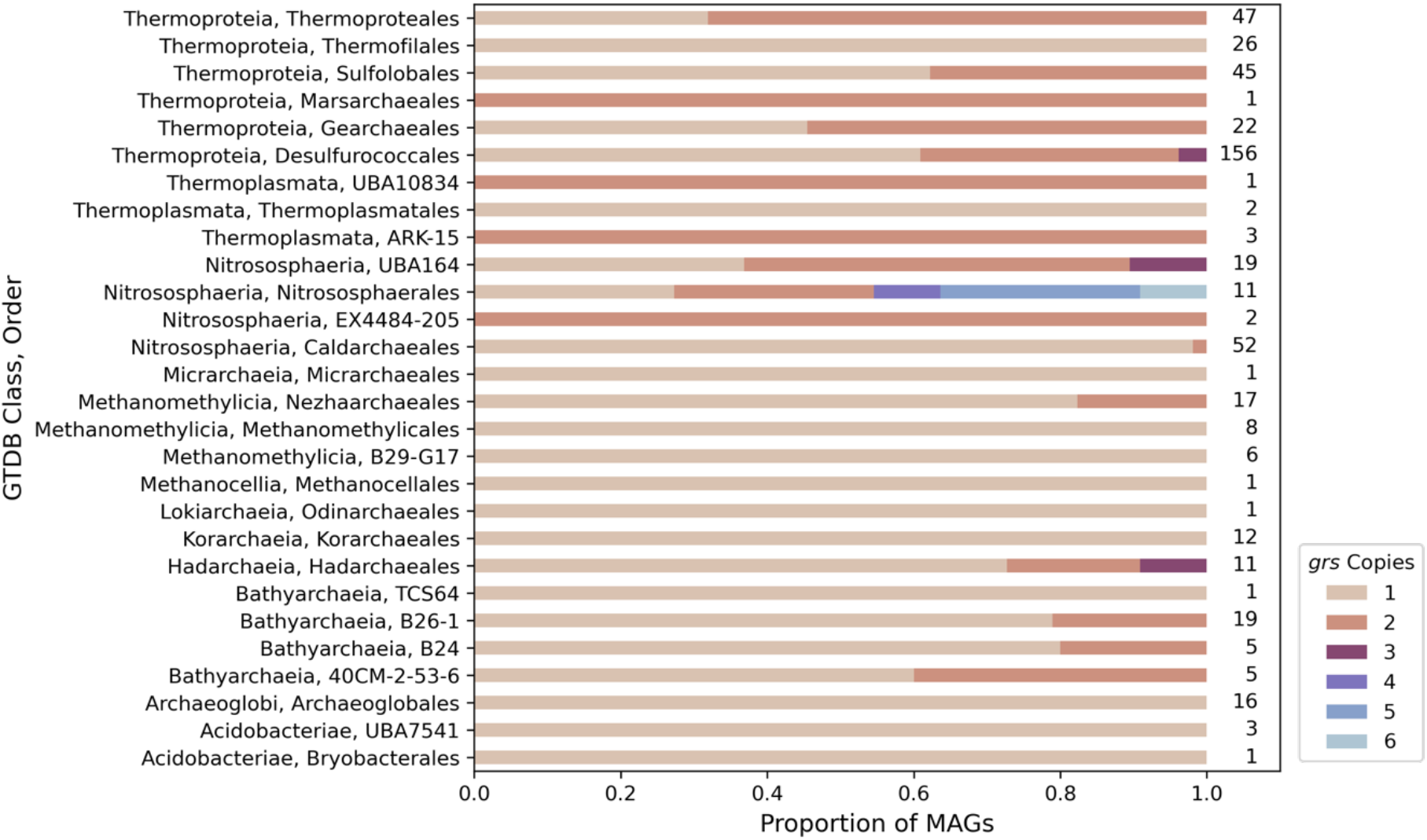
Variation in *grs* copy number across taxonomic orders. The copy number of *grs* varies at the order level in 494 MAGs assembled from thermal springs metagenomes. 494 out of 501 total *grs*-encoding MAGs were classified to the order level. These are the same data shown in Figure 4, but broken down at the order level.

**Figure S5.**
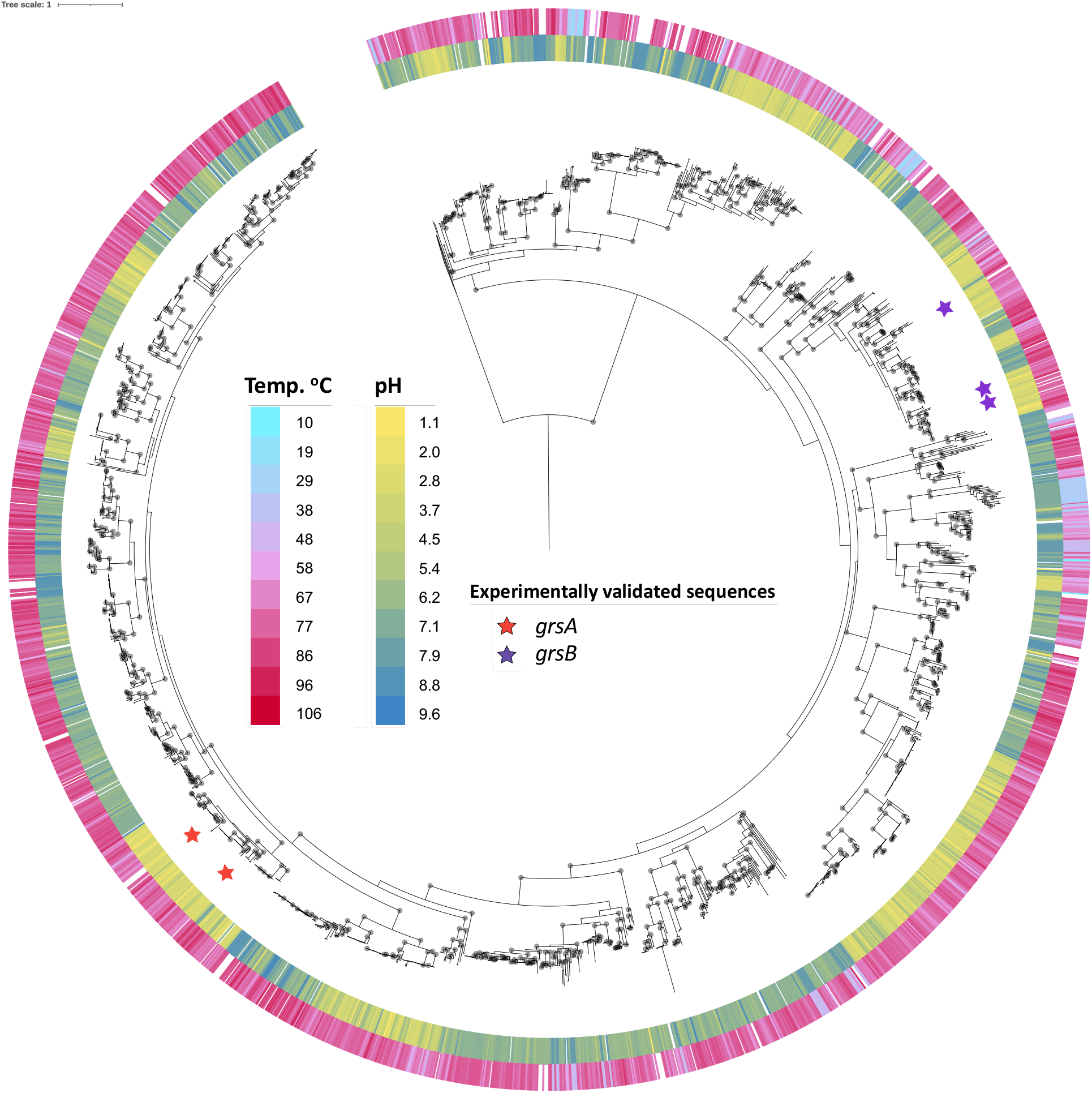
Phylogenetic relationships of the *grsA and grsB* homologs. A rooted maximum likelihood phylogeny of 2,112 *grs* homolog sequences from 307 thermal springs datasets. Sequences are primarily from metagenomes (n = 1,932 sequences) but also include metatranscriptomes (n = 76), cultured isolates (n = 53), metagenome-assembled genomes (n = 47), and single cell genomes (n = 4). Experimentally validated *grsA* and *grsB* sequences (42) are marked by red and purple stars, respectively. Bootstrap support values ≥75% are marked by circles. The outer color bars correspond to sample temperature and the inner color bars correspond to sample pH. Tree scale is in units of substitutions per site.

**Supplementary Tables, available at:** https://doi.org/10.6084/m9.figshare.c.5874383.v1)

**Table S1**. Amino acid reference sequences of *grs* homologs used to build the HMM profile.

**Table S2**. Datasets searched including accession in JGI’s IMG database and associated metadata.

**Table S3**. Breakdown of sample types (e.g. metagenome, isolate genome, single cell analysis, etc.) as cataloged in JGI’s IMG database.

**Table S4**. Geographic locations of (meta)genomic samples as reported in JGI’s IMG database.

**Table S5**. Number of *grs* sequences detected per sample.

**Table S6**. *grs* abundance metrics and geochemistry used for metagenomes in Figure 3, Figure S2, and Figure S3.

**Table S7**. Conversion between GTDB and NCBI taxonomy nomenclature to accompany phylogeny.

**Table S8**. Thermal springs isolates reported in Figure 4, including known lipid profiles, growth pH and temperature.

**Table S9**. MAGs searched in this analysis and *grs* copies detected.

**Table S10**. Summary of taxonomic affiliation of MAGs in which *grs* homologs were detected.

**Table S11**. Annotations for the Figure 5 phylogeny.

**Table S12**. Physicochemical properties of amino acids used in the naïve Bayes classifier to categorize grsA-like sequences. Inspired by Kogay et al., 2019.

**Table S13**. Predictions from the machine learning classifier.

/end

